# A comparison of tools and read depth criteria for genuine single nucleotide polymorphism identification in ancient maize samples

**DOI:** 10.1101/2025.07.22.666096

**Authors:** Huan Chen, Thelma F. Madzima

## Abstract

Maize is an important subject in the study of ancient DNA (aDNA) due to its profound historical, cultural and economic significance. Studies of maize aDNA can provide insights into domestication, evolution and gene flow of this important crop plant. During analysis of aDNA samples, it is essential to elucidate the extent of post-mortem damage (PMD) as well as to select optimal tools and read depth criteria for genuine single nucleotide polymorphism (SNP) discovery. To date, standardized approaches to address these issues are lacking. Using eight archaeological maize samples from publicly available datasets, we compared the performance of two different approaches for solving PMD, Rm5nt and NoChange. We also compared three different analysis tools (GATK, angsd and pileupCaller) and read depth criteria (2X, 5X and 10X) for genuine SNP discovery of aDNA maize samples. The results showed that the Rm5nt approach performed better in processing aDNA. We also found that the angsd tool with a read depth >= 5X is optimal for inferring the closest relative of the aDNA maize samples used in this study. Our study aims to improve the current standardized approach, including optimal approaches, tools and read depth criteria, to improve the accuracy of aDNA data interpretation. Furthermore, our results offer a practical guideline for researchers seeking to improve the quality of their aDNA data for downstream analysis.

## INTRODUCTION

Ancient DNA (aDNA) can have a time depth and age of tens of thousands of years before present (BP), and the genetic information gathered from aDNA is crucial for understanding the evolutionary process of humans, the domestication history of animals and plants, and other species, and understanding the evolutionary trajectory of modern species. For example, aDNA studies of Neanderthals have helped us understand their evolutionary origins, migrations, biological interactions, and way of life, as well as genetic connections between ancestral human population and anatomically modern humans[1, 2]. aDNA studies of crops such as maize have expanded our knowledge of gene flow in North, Middle and South America, and following European contact across the globe, thereby providing insights into the history of domestication, and its influence on modern maize. Since the first aDNA research began in 1984[3, 4], research using aDNA samples has increased, aided by advances in DNA extraction techniques, as well as sequencing and bioinformatics technologies and methods, expanding our knowledge about the past[5]. Next-generation sequencing (NGS) technologies play an essential role in aDNA research advances through the ability to analyze DNA at the single nucleotide polymorphism (SNP) level and shifting the focus from analyzing SNPs from a few known genes to the whole genome. Genuine SNPs, are true SNPs that are biologically validated, rather than the result of sequencing errors or technical artifacts, and are essential for reliable aDNA analysis. For example, using genuine SNPs from individual ancient maize samples, researchers have been able to identify maize domestication and adaptation, evolution and gene flow[6-10].

When analyzing aDNA samples from maize, the common first step is to find the closest relative of the target samples using principal component analysis (PCA). This has appeared to be the preferred approach ever since PCA was first applied to genetic data in 1978[11]. SNPs from whole genome sequencing (WGS) are used for PCA to infer the closest relative for target aDNA samples. Post-mortem damage (PMD) (e.g., cytosine deamination to uracil), that results in high nucleotide misincorporation at sequence read ends, has been suggested as a powerful signature to authenticate aDNA from NGS data[12]. Although approaches and tools have been developed to address the caveat of PMD and data processing methods, the key criteria - such as filter criteria for accurately identifying genuine SNPs that meaningfully contribute to the interpretation of the final results - remains a challenge. This continues to limit the accuracy of genuine SNP discovery and their impact on downstream analysis[12-15].

Various tools and read depth criteria are used for genuine SNP discovery. For example, there are mapping tools such as bwa aln, bwa mem, and SNP discovery tools such as GATK[16], ANGSD[17], PileupCaller[18] as well as set read depth criteria[6-10, 19-23]. These SNPs are subsequently used in PCA and other downstream analysis. However, there is a lack of a “gold standard” of what tools and read depth criteria should be used to identify SNPs that genuinely reveal the story behind aDNA samples in maize studies.

Here, we describe a comparison among three different tools and read depth criteria that are commonly used for the identification of genuine SNPs in aDNA analysis of maize samples. Our results show that the different tools and read depth criteria affect the inference of the closest relative for target aDNA samples.

## METHODS

### Archaeological sample selection

We selected eight archaeological maize samples for our analysis: Arica4, Z2, Z6, Z61, Z64, Z65, Z66, Z67 (Table S1) that are publicly available from previous studies, with age range from 5,000BP to 100BP[6, 7]. We chose these eight samples because their genotypes were determined by both direct genotype calling and pseudohaploid calls, making downstream comparisons more suitable for the purposes of this study.

### Data processing

The sequences of the eight archaeological maize samples were downloaded from the Sequence Read Archive (SRA) (https://www.ncbi.nlm.nih.gov/sra/; Table S1). The raw sequence (SRA format) for each sample was converted into FASTQ format using fastq-dump (https://edwards.flinders.edu.au/fastq-dump/). Adapters were removed and paired reads were merged using AdapterRemoval[24] with parameters -- minquality 20 --minlength 30. Reads from all samples were solved using two different approaches. In the first approach, all 5′ thymine (T) and 3′ adenine (A) residues within 5 nucleotides (5nt) of the two ends were hard-masked as these are loci where deamination was is most concentrated (T and A -> N; where N represents unknown types of nucleotides)[6, 7]. We named this processing approach the “Remove 5nt” (Rm5nt) group, as previously described by Kistler, L. *et al*[6, 7]. For the second approach, we retained reads that did not exhibit any change at 5′ thymine (T) and 3′ adenine (A) residues within 5 nucleotides (5nt) of the two ends were kept as they are. We named this processing approach the “NoChange” group. For both approaches, reads were then mapped to the soft-masked B73 version 5 reference genome using the Burrows-Wheeler Alignment (BWA) tool v0.7.17 with disabled seed (−l 1024 -o 0 -E 3) and a quality control threshold (−q 20) based on the recommended parameter[25] to improve ancient DNA mapping. Duplicated reads were removed by Picard (https://gatk.broadinstitute.org/hc/en-us). The read depth of each nucleotide position was estimated by bam-readcount (https://github.com/genome/bam-readcount).

### Variant calling

We used three different tools with different algorithms for SNP discovery in the archaeological maize samples. 1) GATK[16] was used to directly generate genotype calls. The commands executed were: gatk --java-options “-Xmx4g -Xms4g” GenotypeGVCFs -reference Reference.fa -V gendb:/path/genomicsdb -output output.vcf. 2) The angsd[17] tool was used to generate pseudohaploid calls. The commands executed were: angsd -bam <ancient sample name list> -dohaplocall 1 -doCounts 1 - nThreads 16 -out output. 3) SequenceTools pileupCaller v1.5.4.0[18] allowed us to generate pseudohaploid calls for the eight archaeological maize samples. The commands executed were: samtools mpileup -R -B -q20 -Q20 & pileupCaller –sampleNames <ancient sample name> randomHaploid --singleStrandMode. These three processing tools yielded SNPs for both the NoChange and Rm5nt approaches.

As several SNP loci had more than four types of nucleotide bases (e.g., A,G,C,T,N) in pseudohaploid calls using angsd[17], these SNPs were removed from the analysis because they stopped converting haplo into vcf files. We further filtered the SNPs using estimated information from bam-readcount (https://github.com/genome/bam-readcount). SNPs supported by at least two, five and ten reads (Read Depth >= 2X, 5X and 10X) were retained for each individual ancient maize sample. We then combined the filtered SNPs from each sample, generated through different tools and read depth criteria, for subsequent PCA analysis.

## RESULTS

### The Rm5nt approach increases mapping accuracy

The estimated results from mapDamage show high levels of deamination at residues within 5nt of the 5’ and 3’ ends in aDNA samples[26]. Therefore, we used two alternative approaches: the Rm5nt and NoChange. To determine which processing approach would be less sensitive to PMD and generate more accurate and genuine SNPs for analysis, we compared the Rm5nt and NoChange data groups. Total read counts were variable between the eight samples (Figure 1A and Dataset S1), although we consistently observed that the samples in the Rm5nt group had higher read counts than the NoChange group after removing adapters and merging paired reads, and that the total read count that passed quality control (QC-passed) were also higher when compared to the NoChange groups. However, the mapped read counts of the NoChange group were 3.8 to 5.2 times higher than the Rm5nt group (Figure 1B). A similar observation was observed in the mapping percentage where the NoChange group values were higher than the Rm5nt group. Interestingly, the mapping percentages were found to be very diverse among these eight samples, with variations ranging from 6.82% to 95.15% and 0.34% to 22.18%, in the NoChange and Rm5nt groups, respectively (Figure 1C). These findings indicate that the Rm5nt approach could increase the read count and quality for merged paired reads, but the mapping rate is lower when compared to using the NoChange approach. These changes are possibly due to the fact that the Rm5nt approach increases the mapping accuracy by hard-masking the potential PMD bases within 5nt that were used to align to the reference genome using bwa aln with seed disabled mapping.

**Figure 1.**
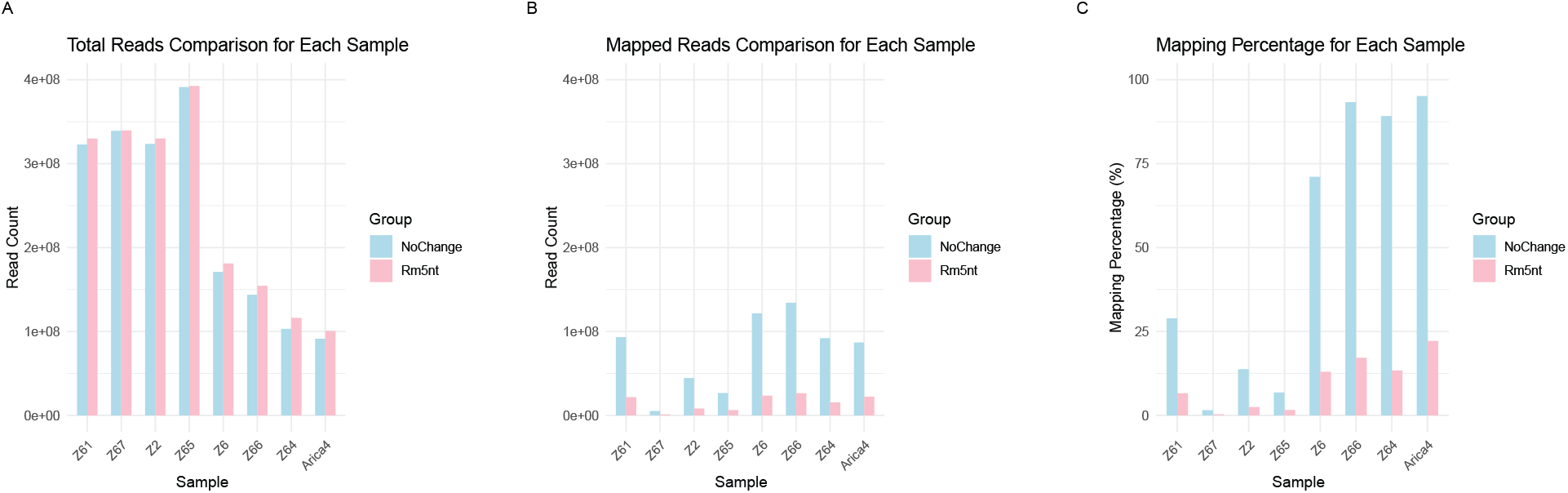
Read count statistics. A. Total read count, B. Mapped reads, C. Mapping percentage for each sample between NoChange and Rm5nt groups. Blue and pink indicate NoChange and Rm5nt groups, respectively.

### The Rm5nt and NoChange approaches perform differently in SNP discovery

SNP discovery in aDNA presents unique challenges due to PMD, and low sequencing depth. Therefore, appropriate tools for genuine SNP discovery are critical to correctly and accurately interpret results. In this study, we explored three different tools for SNP discovery. The results showed that the SNP count is the highest using angsd[17], while very low using GATK[16] in the Nochange group (Figure 2A and Dataset S2). Although in the Rm5nt group, the SNP count using pileupCaller [18] is the highest, but considering ∼40% SNPs are missing in all samples, angsd[17] still has the highest count for SNP discovery (Figure 2B and Dataset S2). Similar patterns were observed across all ten chromosomes.

**Figure 2.**
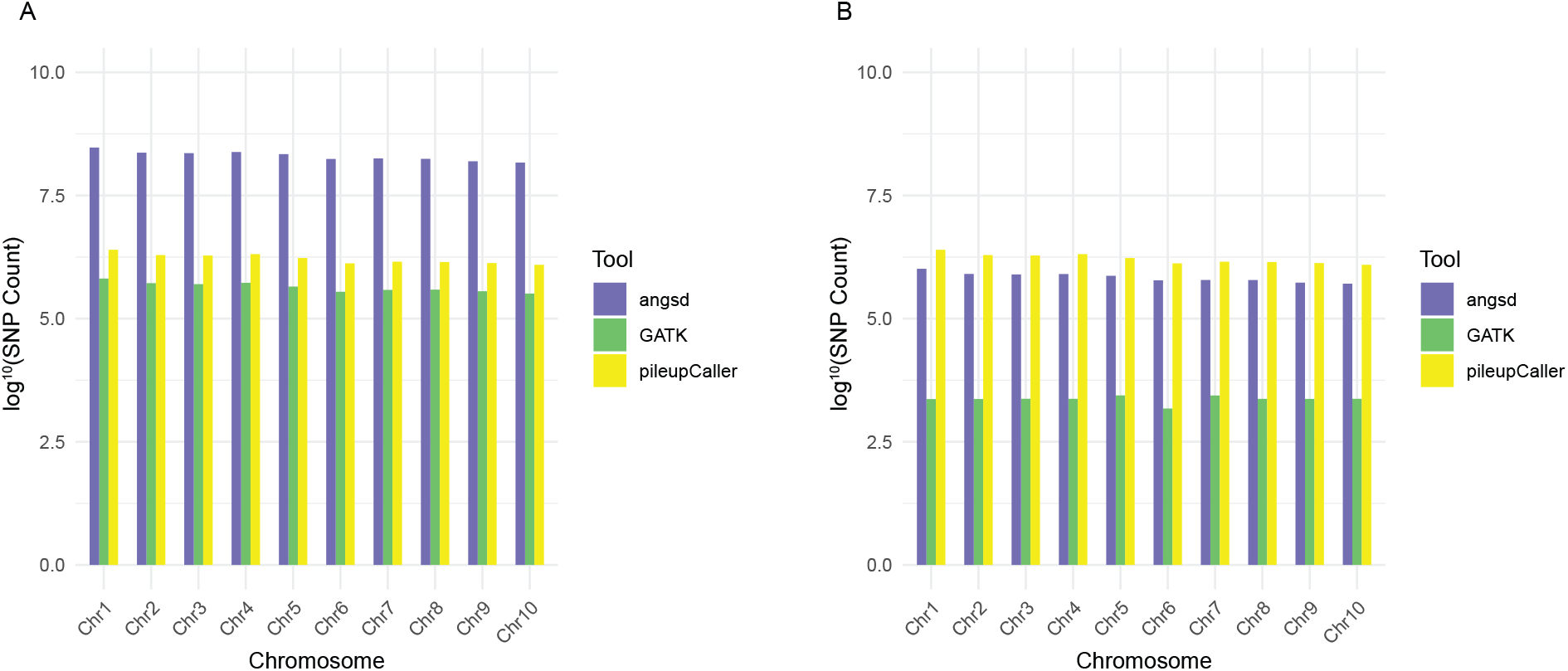
Single nucleotide polymorphism (SNP) count statistics. A. log10 SNP count in the NoChange group. B. log10 SNP count in the Rm5nt group. Blue and pink indicate NoChange and Rm5nt groups, respectively.

Sequencing read depth is one of the key factors for genuine SNPs discovery, and the accuracy of SNPs detection increases with read depth. However, due to damage to the aDNA genome and the needed balance between sequence depth and cost, the read depths (e.g., 10 - 30X) are not always stringent enough for accurate identification of SNPs. We were interested in determining how the accuracy of SNP detection would be affected by read depth and the impact on overall interpretation of aDNA research results. To address this question, we calculated the read depth for each nucleotide (Figure 3 and Dataset S3). SNP counts for read depths equal to or higher than 2, 5 and 10X coverage show that the NoChange group had higher nucleotide loci count than the Rm5nt group (Figure 3). In both groups, the nucleotide location counts decreased following read depth increase for all the samples. Interestingly, the location counts are very close at 10X read depth, although the numbers vary at 2X among all samples. Compared to the 2X read depth, 4.45% to 41.23% and 2.37% to 17.57% of nucleotides remained at a 5X read depth in both the NoChange and Rm5nt groups, respectively. For the 10X read depth, 0.47% to 6.10% of nucleotides remained in the NoChange group, but only 0.15% to 1.68% of nucleotides remained in the Rm5nt group (Figure 3 and Dataset S3). This indicates that only 2% or fewer nucleotides with 10X read depth meet the criteria to appear in the Rm5nt group. This led us to question whether this stringent threshold of 2% accurately represents aDNA data for downstream analysis and interpretation of results from procedures such as PCA.

**Figure 3.**
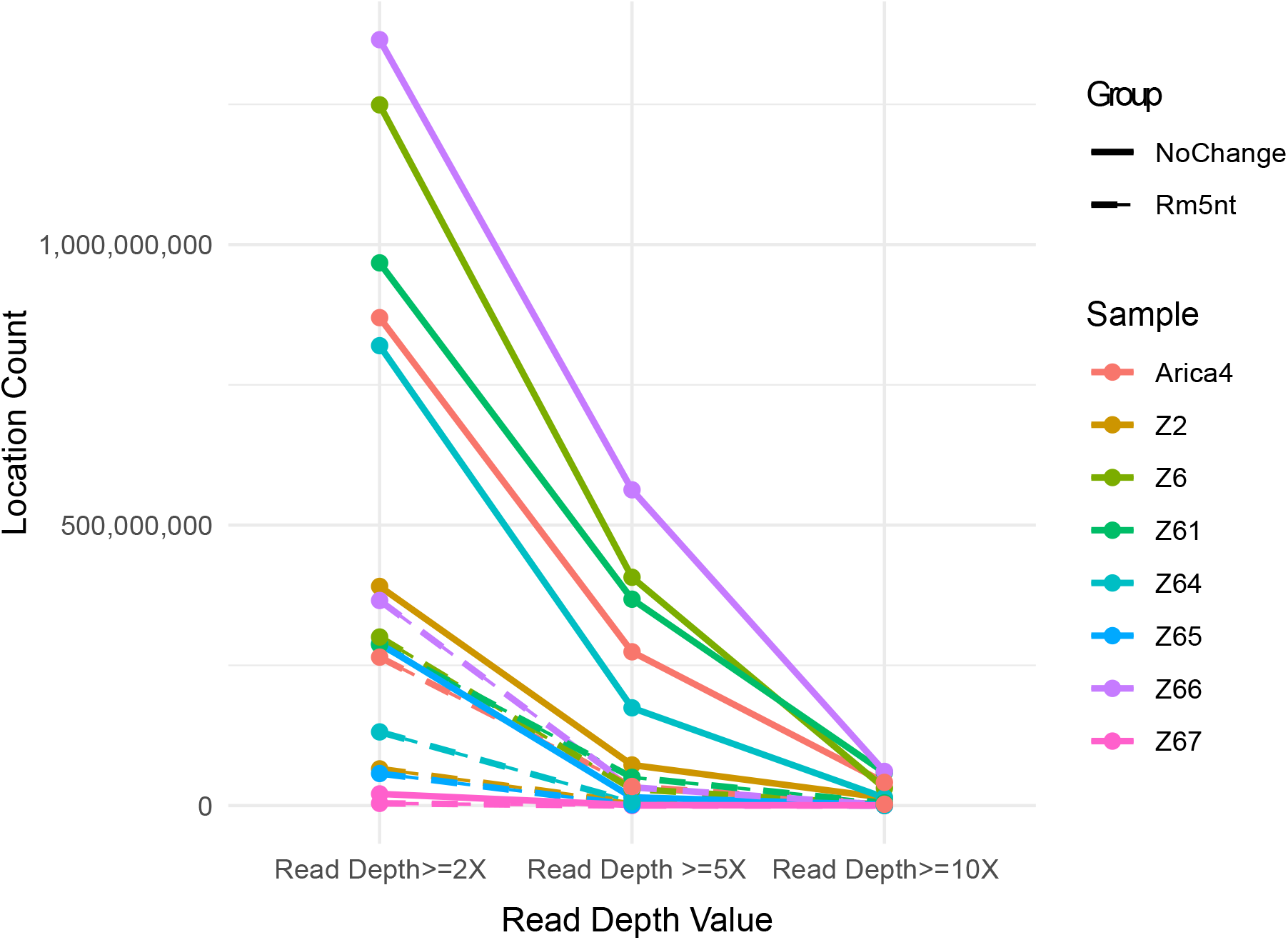
Read depth value distribution. Different color indicates different samples. Solid and dotted lines indicate NoChange and Rm5nt groups, respectively.

Combining the above information, we calculated the SNP counts discovered by the three tools at three different read depth criteria by checking how different combinations of tools and read depth criteria affect the SNP counts. The results showed that angsd[17] tool detected the highest number of SNPs, followed by pileupCaller[18], which detected the second-highest counts, while GATK[16] detected the fewest SNPs at 2X, 5X and 10X read depths (Figure 4 and Dataset S4). Interestingly, while the SNP counts differ between these tools, the overall trends in change are similar, where counts decrease from 2X to 5X to 10X across all samples. However, tool bias appears to affect the SNP discovery particularly in sample Z67 due to it being only ∼100 years BP.

**Figure 4.**
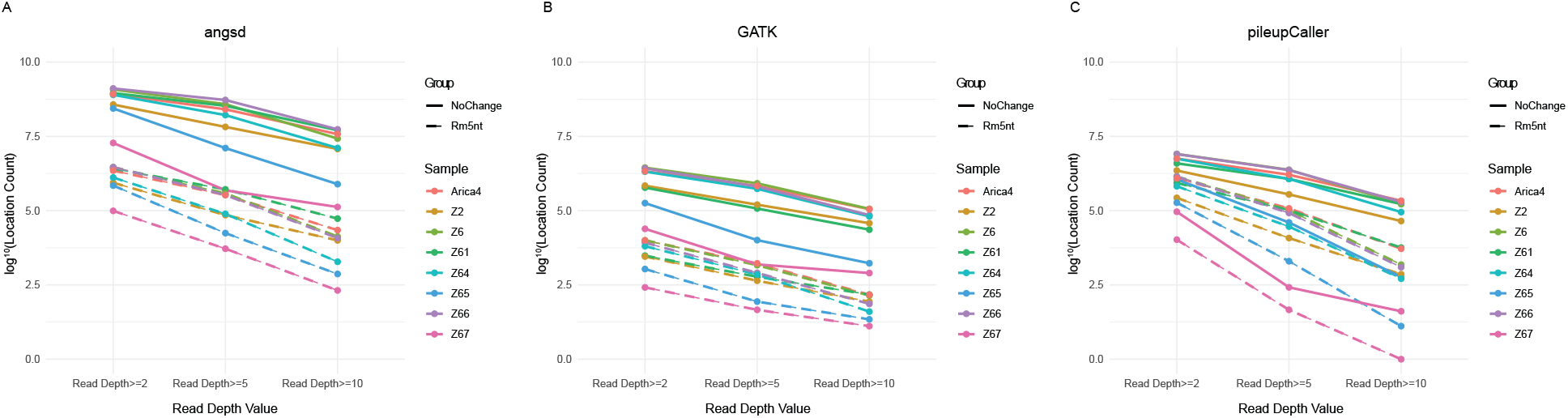
Single nucleotide polymorphism (SNP) count statistics with different read depth criterions. Different colors indicate different samples. Solid and dotted line indicate NoChange and Rm5nt groups, respectively. Depth >= 2X, 5X and10X indicates at least 2, 5, 10X reads are mapped on this SNPs.

### The Rm5nt approach is the better one to process aDNA data

We were curious about the SNPs that were discovered by the three tools: GATK[16], angsd[17] and pileupCaller[18]. We found that the count of overlapping SNPs that share the same location on the genome among these three tools decreased as the read depth criteria increased, and the overlapping SNPs when using the Rm5nt approach were much lower than the NoChange data (Figure 5 and Dataset S5). This indicates different tools have their own bias in SNP discovery for aDNA maize samples.

**Figure 5.**
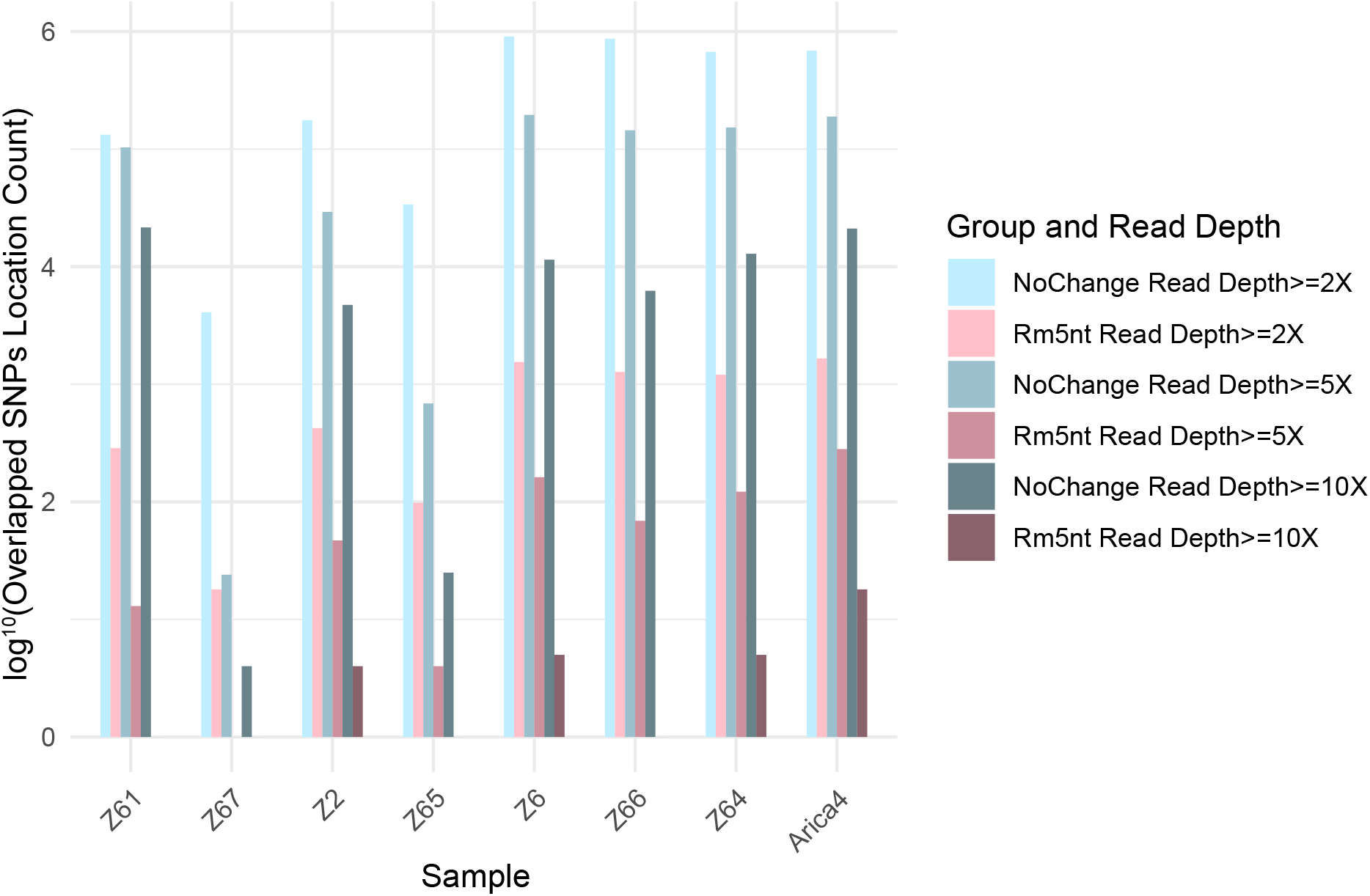
The overlapped single nucleotide polymorphism (SNP) among angsd, GATK and pileupCaller with different read depth criterions. Blue and pink indicate NoChange and Rm5nt groups, respectively. Depth >= 2X, 5X and 10X indicates at least 2, 5, 10X reads are mapped on these SNPs.

GATK[16] is widely used for SNP discovery and has been applied to ancient maize samples. However, our results show that it may not perform well on low-coverage data because it is not tolerant of low read depth and its hard filters and default settings may exclude valid but low-confidence SNPs. Considering it is not good for SNPs discovery with low read depth data, we proceeded to only focus on the overlapping SNPs detected using angsd[17] and pileupCaller[18] that are commonly used to generate pseudohaploid calls for aDNA in particular. The overlapping SNP counts from the Rm5nt group was much lower than the NoChange group across different read depth criteria (Figure 6A and Dataset S6). We also calculated the SNPs, which not only share the same location on the genome, but also have the same reference allele (REF) and alternative allele (ALT) information. Convincingly, the NoChange group still had higher counts than Rm5nt group, but surprisingly, the ratios of overlapping SNPs with the same REF and ALT information in Rm5nt, which are 3.1% to 13.9, 6.8% to 50.0%, 13.1% to 30.4% at 2X, 5X and 10X, respectively, were higher than NoChange group, which are 1.8% to 4.8, 2.0% to 10.2%, 2.0% to 11.9% at >=2X, 5X and 10X, respectively (Figure 6B and Dataset S7). This indicates the Rm5nt approach performs better when processing aDNA maize samples. But this also raises our concerns about how accurate the SNPs could be, and how strong the bias is when using different tools and read depth criteria for SNP discovery of aDNA maize samples.

**Figure 6.**
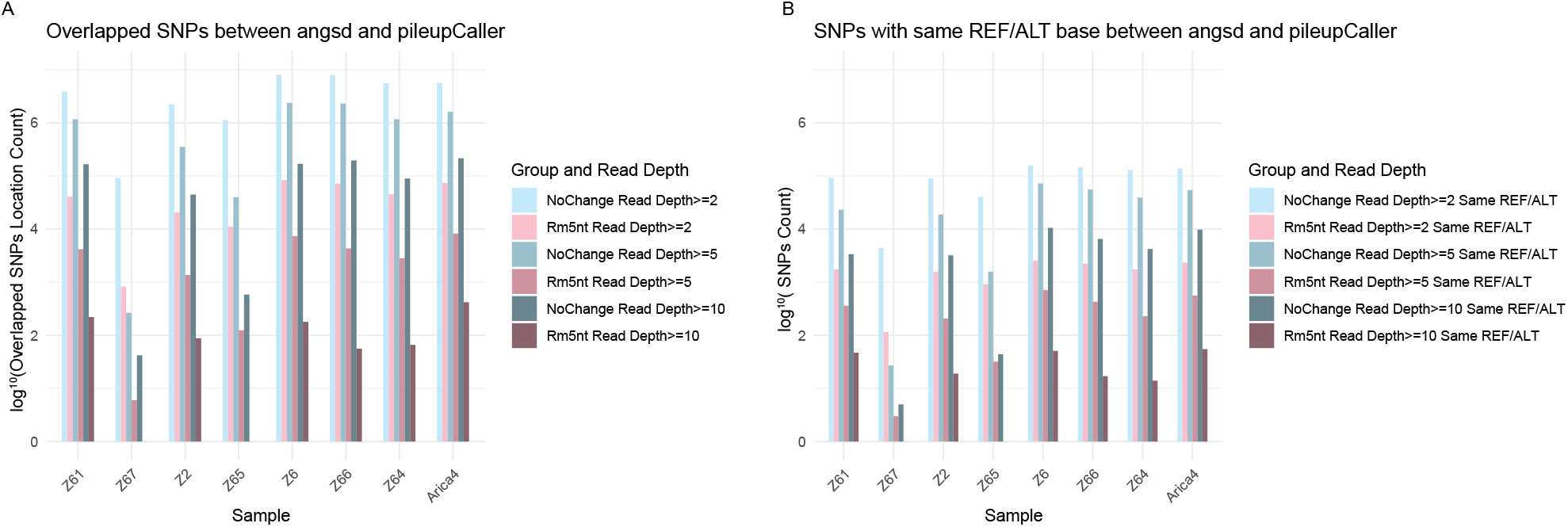
The overlapped single nucleotide polymorphism (SNP) between angsd and pileupCaller with different read depth criterions. A. The overlapped SNPs between angsd and pileupCaller. B. The overlapped SNPs that have the same REF and ALT information between angsd and pileupCaller. Blue and pink indicate NoChange and Rm5nt groups, respectively. Depth >= 2X, 5X and 10X indicates at least 2, 5, 10X reads are mapped on these SNPs.

### Rm5nt approach with angsd at 5X is the better one to infer the closest relative

The above results show that the Rm5nt approach performs better in the analysis of aDNA maize samples. However, maintaining a good balance between SNP counts and read depth makes it difficult to find an optimal solution that will produce not only an adequate number of SNPs but also genuine SNPs for downstream analysis, due to the lack of a “gold standard”. To address this challenge, we applied PCA for the SNPs, with different combinations of various tools and read depth criteria, to infer their closest relative, which potentially helped us identify the best criteria for maintaining a good balance.

The PCA results for SNPs from angsd[17] showed that sample similarity ratio based on the first two components increased from 41.2% to 42.9% and 53.0% when the read depth threshold was raised from >=2X, 5X and 10X, respectively. However, the distances between samples are changed across various read depth criteria (Figure 7). Notably, samples Z65 and Z67 exhibited the longest distance to all other samples at read depth >= 2X and 5X, but not at >=10X. Interestingly, the distances among all samples – excluding Z67-increased as read depth increased. Across all PCAs, sample Z67 consistently showed the farthest distance from all the other samples. PCA results based on SNPs from both GATK[16] and pileupCaller[18] at read depth >= 5X and 10X were excluded due to their insufficient data to meet requirements for the analysis (Figure 7). The results indicate that both the clustering pattern and distances among samples differ from those observed with angsd[17].

**Figure 7.**
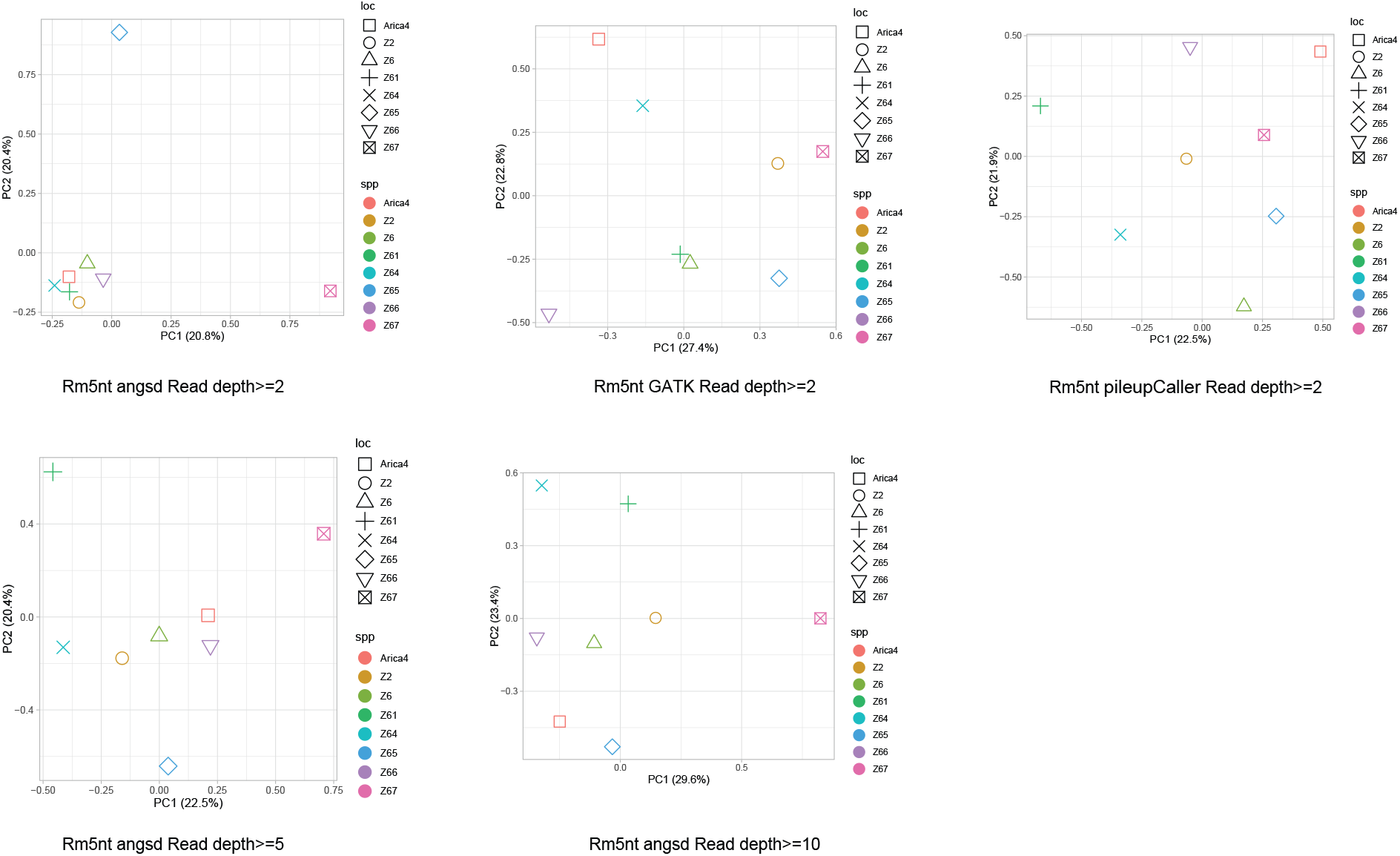
Principal component analysis (PCA) of nuclear genomic data from aDNA maize. Samples with greater affinity to one another cluster closer together.

In summary, the conclusions based on different combinations of tools and read depth criteria lack robustness. To have a standard for identifying the best criteria, we used distance in PCA analysis from previous research as our reference[7]. The results indicated that PCA from Rm5nt with read depth >= 5X generated by angsd[17] has the most similar pattern as the reference. This suggested that the Rm5nt approach with read depth >= 5X using angsd[17] provides the best criteria for keeping a good balance between SNP counts and read depth in our analysis, and it may be suitable for analyzing other aDNA maize samples.

## DISCUSSION AND CONCLUSION

Improving the accuracy of genuine SNP discovery from aDNA that contribute to downstream analysis is essential to accurately and correctly uncovering the story behind archeological genetic information in maize, especially for an understanding of maize evolution. However, the best methods to improve the accuracy of genuine SNP identification for analysis is not well described. Reducing the impact of PMD on downstream analysis is one aspect where we can work to improve the analytic accuracy[12]. Our results using two different approaches for processing data suggest that hard-masking the PMD nucleotide base within 5nt (Rm5nt) is essential for improving the accuracy of SNP discovery, and this is also consistent with approaches that were used in previous studies[6, 7, 9, 10]. However, the Rm5nt approach is not the best for analysis of all aDNA maize samples due to sample variability. Damage estimate tools, such as mapDamage2[26], need to be used first to estimate misincorporation frequencies for each sample, before applying hard-masking to the nucleotide bases where deamination was most concentrated. As such, the hard-masked values will vary depending on different aDNA samples.

Choosing optimal tools for genuine SNP discovery is another aspect that we can work on to improve the analysis accuracy. Our results showed that three tools and three read depth criteria suggest that angsd[17] performs better than other tools in SNP discovery. However, one thing left to consider is that angsd[17] will exclude a small part of the SNP loci that have more than four types of nucleotide bases, such as A,G,C,T,N. For example, as shown in the results, 45 SNPs were removed from the Rm5nt group, while 6,452 SNPs were removed from the NoChange group. This raises the question that whether this small portion SNP is essentially important or key SNPs for aDNA studies. If so, it may cause the loss of valuable information for aDNA analysis, and might mislead the concluding interpretation. This issue will be more apparent when a large proportion of samples have low sequencing depth, and more SNPs will be removed due to SNP loci that contain more than four types of nucleotide bases.

Read depth is also an aspect that warrant further work to improve analytic accuracy. Our results from three read depth criteria suggest that read depth >= 5X may be a good balance between SNPs count and read depth for aDNA analysis (Figure7). However, it also depends on the specific question that was asked for the specific aDNA analysis. For example, if we only focus on the question of whether the Z67 sample is more distant from all other samples, then read depth >= 2X, 5X and 10X are all equally good criteria since they generated the same result (Figure7).

In summary, due to the small ancient sample sizes in our study, our conclusions may have certain limitation in their applicability to all aDNA maize studies. However, the results demonstrate that benchmarking tools with optimal criteria and approaches is critical for effectively processing aDNA maize samples. Our results suggest that, for aDNA maize analysis 1) hard-masking the PMD nucleotide base is essential; 2) GATK[16] is not a good tool for SNPs discovery; 3) angsd[17] has the better performance; and 4) a good balance between SNP count and read depth criteria is important for downstream analysis. Collectively, our approach may help improve the accuracy of genuine SNP discovery during analysis of aDNA samples in maize.

## ACKNOWLEDGMENTS

We deeply thank Dr. William Lovis, Michigan State University, for providing critical feedback for this research. This work is funded by Michigan State University start-up funds to TFM.

## Conflict of Interest

none declared.

**Table S1.**
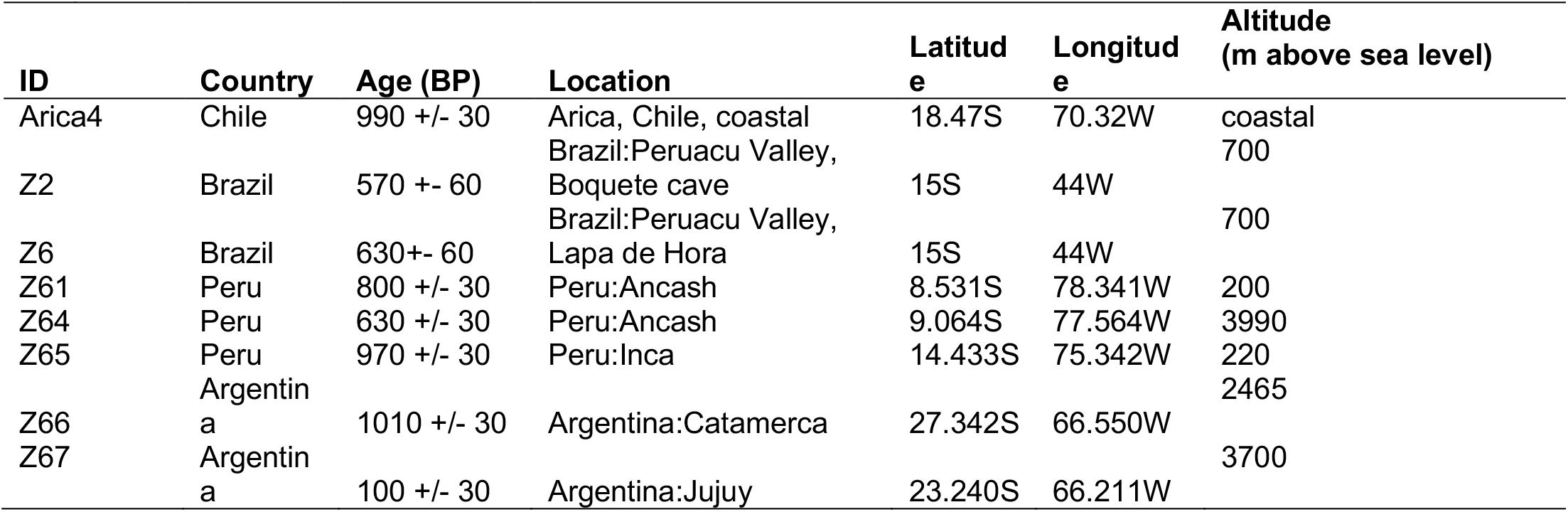
Geographical information of the archaeological maize samples used in this study.

